# Predicting the impact of pneumococcal conjugate vaccine programme options in Vietnam: a dynamic transmission model

**DOI:** 10.1101/121640

**Authors:** Olivier Le Polain De Waroux, W. John Edmunds, Kensuke Takahashi, Koya Ariyoshi, E. Kim Mulholland, David Goldblatt, Yoon Hong Choi, Duc-Anh Dang, Lay Myint Yoshida, Stefan Flasche

## Abstract

**Background:** Catch-up campaigns (CCs) at the introduction of the pneumococcal conjugate vaccines (PCVs) may accelerate the impact of PCVs. However, limited vaccine supplies may delay vaccine introduction if additional doses are needed for such campaigns. We studied the relative impact of introducing PCV13 with and without catch-up campaign, and the implications of potential introduction delays.

**Methods:** We used a dynamic transmission model applied to the population of Nha Trang in Sout central Vietnam. Four strategies were considered: routine vaccination (RV) only, and RV alongside catch-up campaigns among <1y olds (CC1), <2y olds (CC2) and <5y olds (CC5). The model was parameterised with local data on human social contact rates, and was fitted to local carriage data. Post-PCV predictions were based on best estimates of parameters governing post-PCV dynamics, including serotype competition, vaccine efficacy and duration of protection.

**Results:** Our model predicts elimination of vaccine-type (VT) carriage across all age groups within 10 years of introduction in all scenarios with near-complete replacement by non-VT. Most of the benefit of CCs is predicted to occur within the first 3 years after introduction, with the highest impact in the first year, when IPD incidence is predicted to be 11% (95%CrI 9 – 14%) lower than RV with CC1, 25% (21 – 30 %) lower with CC2 and 38% (32 – 46%) lower with CC5.

However, CCs would only prevent more cases of IPD insofar such campaigns do not delay introduction by more than 31 (95%CrI 30 – 32) weeks with CC1, 58 (53 – 63) weeks with CC2 and 89 (78 – 101) weeks for CC5.

**Conclusion:** CCs are predicted to offer a substantial additional reduction in pneumococcal disease burden over RV alone, if their implementation does not result in much introduction delay. Those findings are important to help guide vaccine introduction in countries that have not yet introduced PCV, particularly in Asia.

## Background

Disease due to *Streptococcus pneumoniae* (the pneumococcus) is a leading cause of morbidity and mortality worldwide, disproportionally so in resource-poor settings [1-3].

Ten- and 13-valent pneumococcal conjugate vaccines (PCV10 and PCV13), which cover 10 and 13 of 94 known serotypes [4], are steadily being introduced into the routine immunisation programmes of many low and lower-middle income countries with the support from Gavi, the Vaccine Alliance [5]. In 2015, 50 out of 73 countries eligible for Gavi support had introduced the vaccine, and eight additional countries have been approved for introduction and are expected to introduce PCV within the next two years [5]. In 2012 Pakistan was the first Asian country to routinely introduce PCV, followed by Nepal, Cambodia and Lao PDR [5]. However, PCV has not been introduced in most of Asia, including Vietnam. Globally, the number of infants who had had 3-doses of PCV remains low [6].

The WHO recommends introducing PCV into childhood immunisation programmes alongside a catch-up campaign (CC) among older children [7], in order to provide direct protection to age groups at particular risk of pneumococcal disease, as well as to accelerate the population impact of the vaccine through herd protection [8].

However, the magnitude of the additional impact of different CCs over RV remains unclear. Moreover, Gavi has so far not been able to support CCs in eligible countries over concerns that they would increase vaccine introduction delays, given supply constraints [5].

The aim of our study was to explore the differential impact on carriage and invasive disease of catch-up campaigns targeting various age groups, through a dynamic compartmental model of disease transmission, and also explored the possible impact of delaying vaccine introduction to allow for CCs to be undertaken.

## METHODS

The model was applied to the population of Nha Trang (∼360,000 inhabitants), an urban and semi-rural area in south-central Vietnam.

### Data

#### Nasopharyngeal carriage

Two surveys were conducted six month apart among children <5 years randomly drawn from two communes, with 350 children included in January 2008 [9] and another 350 children in July 2008 [10].

Samples were processed and cultured as per WHO recommendations [11]. Serotyping was done by PCR with 29 specific primer pairs which did not differentiate between serotypes 6A and 6B [9]. Given that both antigens are included in PCV13 but that PCV10 does not include 6A, we decided to implement our model for PCV13.

The carriage prevalence of VT and NVT among 5-17 year olds and adults (≥18 years) was estimated based on the prevalence and serotype distribution in children <5 years of age, using a meta-regression model [12].

#### Social mixing patterns

We derived the age-specific contact patterns from a survey conducted in the same area in 2010. The survey included a random sample of 2002 individuals drawn from the population census, who completed a diary similar to the one used in the POLYMOD surveys [13], asking about the frequency and characteristics of social encounters over 24 hours. The corresponding mixing matrix was derived as in Melegaro et al. [14]. We parameterised our model based on physical (i.e. skin-to-skin) contacts only, given that *S.pneumoniae* is generally assumed to be transmitted through close interpersonal contact [15].

### Model structure

We built an age-structured deterministic Susceptible-Infected-Susceptible transmission model of carriage acquisition and clearance, in which we modelled VT jointly and separately from NVT, but allowed for co-colonisation, as in previous models [16]. Details about the model structure and model equations can be found in Supplementary File S1.

The model comprised of three levels of vaccine-induced immunity; (1) no protection, (2) partial protection and (3) full protection. The latter refers to the efficacy and duration of protection conferred after completion of the routine schedule (i.e. 2 infant doses and a booster at 12 months (‘2+1’ schedule)) or the completion of a catch-up programme in older children (2 doses in <18 months and 1 dose in ≥18 months). Partial protection was gained form two primary infant doses, or after the first catch-up dose in children aged 12 - 17 months. The difference between full and partial protection lied in the magnitude of vaccine efficacy against carriage (*VE_C_*) and in the duration of protection.

We applied the model to a population of 81 annual age cohorts (0 to 80 years) divided into 52 weekly age bands of 100 individuals. In the calculation of the force of infection the population figure was adjusted to represent the actual population, based on census data.

### Model fitting

We fitted the model to pre-vaccination nasopharyngeal prevalence data using a Markov Chain Monte-Carlo (MCMC) algorithm, and estimating the age-specific probability of effective transmission in the absence of PCV. For each posterior sample we simulated up to 15 years after vaccine introduction.

In the calculation of the log-likelihood for carriage, model estimates of VT carriage prevalence included VT carriers and VT-NVT co-colonization given the higher likelihood of our PCR assay to detect VT than NVT colonies in case of multiple colonization [9].

### Model parameters and outputs

Table 1 displays the value assigned to the parameters governing transmission and vaccination. Uncertainty around parameters was taken into account by sampling from their posterior distribution in the MCMC process.

**Table 1:**
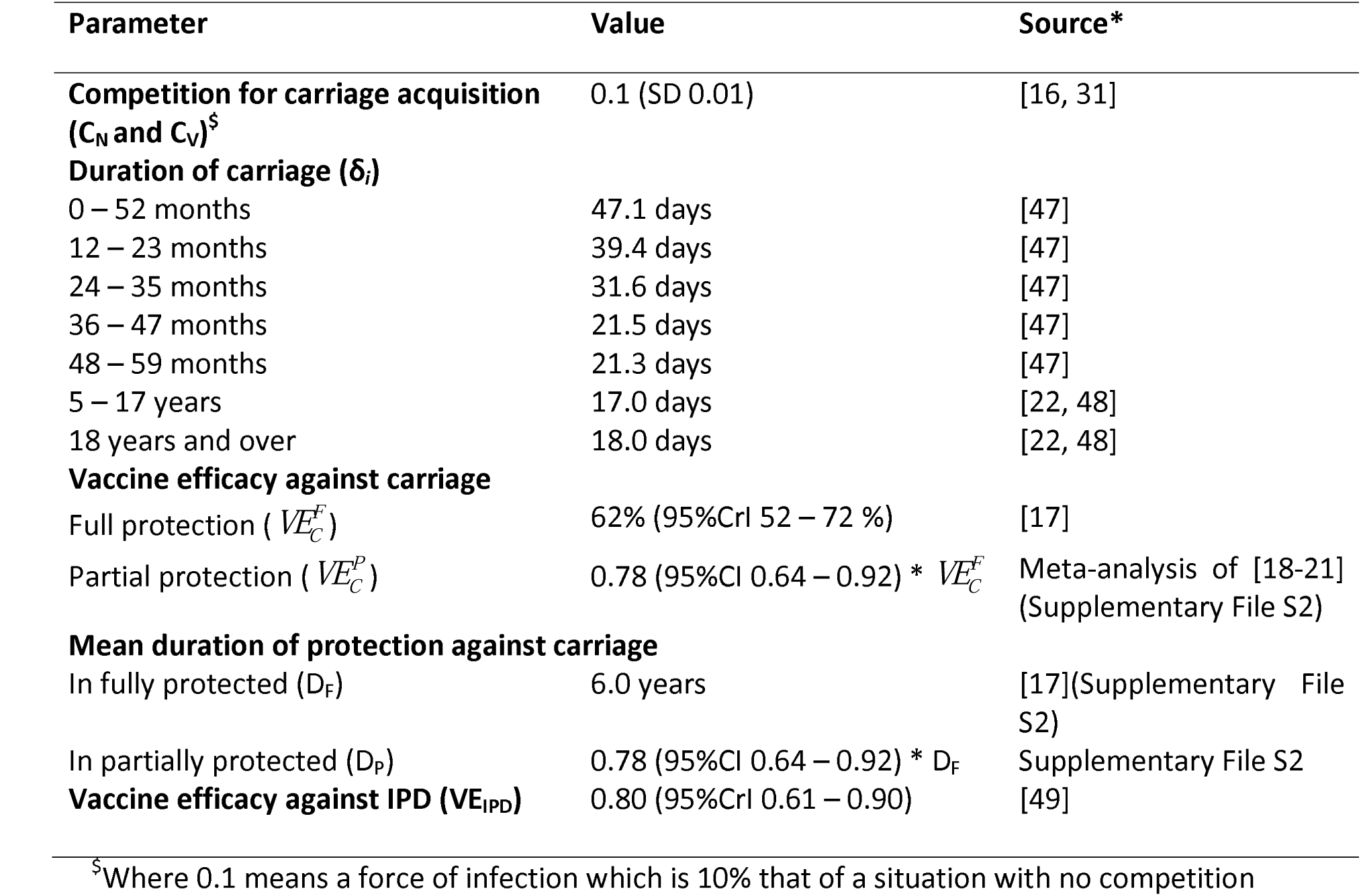
Parameters used in the model.

We obtained the vaccine efficacy against carriage conferring full protection (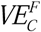) from a meta-regression model [17]. We estimated the partial efficacy against carriage (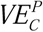) as 0.78 (95% CI 0.64 – 0.92) that of 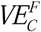 through a meta-analysis of the relative risk of VT carriage after full schedules (2+1 or 3+0) compared to partial schedules (2+0), which included four trials [18-21]. Further details are provided in the Supplementary File S2.

We assumed an exponential decay function for the waning of 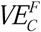, as in previous models [16, 22]. We fixed the average duration of protection to 6 years, which best matched the output of a meta-regression model of waning efficacy [17], and analysed the impact of shorter (3 years) and longer (20 years) average protection in sensitivity analyses, based on the uncertainty bounds of the same model [17]. The average duration of protection following partial vaccination was assumed to be 0.78 (95%CI 0.64 – 0.92) that of a complete schedule, as for efficacy. Details are provided in the Supplementary File S2.

In our base case model we set the vaccination coverage of both routine and catch-up strategies at 90%, in accordance with coverage data from Vietnam [23].

Carriage estimates were translated into IPD estimates based on the propensity of VT and NVT serotypes to cause illness as a result of carriage, by age, using estimates from Choi et al. [24].

The impact of PCV on IPD was calculated based on VT and NVT disease propensity, as well as the efficacy against progression to invasive disease as a result of carriage (VE_inv_). The latter was obtained from a function linking efficacy against IPD (VE_IPD_) with VE_C_ and VE_inv_, where VE_IPD_ =1 − (1- VE_C_) *(1-VE_inv_) [25]. VE_INV_ was calculated based on estimates of VE_IPD_ from Lucero et al. [26] and estimates of VE_C_ derived from [17] (Table1), as detailed in the Supplementary File S2.

Given limited data on IPD from Nha Trang and Vietnam [27], our main analysis focused on the proportion reduction in IPD, rather than IPD incidence. However, we illustrated how the relative IPD change may translate into disease incidence based on a point estimate in <5y olds of 49 /100,000 as reported for Nha Trang [27]. We did not infer disease impact among ≥5 year olds.

To explore the impact of delayed vaccination, we assumed that for each additional week of delay the incidence of IPD during that time would be at its pre-PCV steady level.

### Vaccination strategies

We explored four different strategies: (i) routine (2+1) vaccination only (RV), and routine vaccination with a catch-up campaign in (ii) <1y olds (CC1), (iii) <2y olds (CC2), and (iv) <5y olds (CC5).

WHO currently recommends introducing PCV either as a three primary infant dose schedule (3+0) or as two primary doses with a booster at 12 months of age (2+1 schedule), with the choice between schedules guided by setting-specific epidemiological characteristics [7]. We here present the predicted impact of a 2+1 programme, given the relatively low prevalence of carriage in young children in Nha Trang [27].

### Sensitivity analyses

We also ran the model for coverage levels of 50% and 70% in both routine and catch-up programmes. We also explored the impact of duration of protection on our model outputs, based on lower values of 3 years and 20 years, which span across the range of likely values (Figure S3 in File S2).

## RESULTS

The carriage prevalence in <5 year olds was 41% (95% CI 38 - 46%) overall, 27% (95%CI 23 - 32%) for VT serotypes and 14% (95%CI 11 - 18%) for NVT serotypes. We estimated the carriage prevalence of VT in 5 – 17 year olds and in adults to be 14% (95% CrI 10 – 18%) and 3% (95%CrI 0 – 7%) respectively, and that of NVT to be 15% (95%CrI 11 – 19%) and 3% (95%CrI 0 – 7%). Figure 1 shows the model fit to the carriage data.

**Figure 1:**
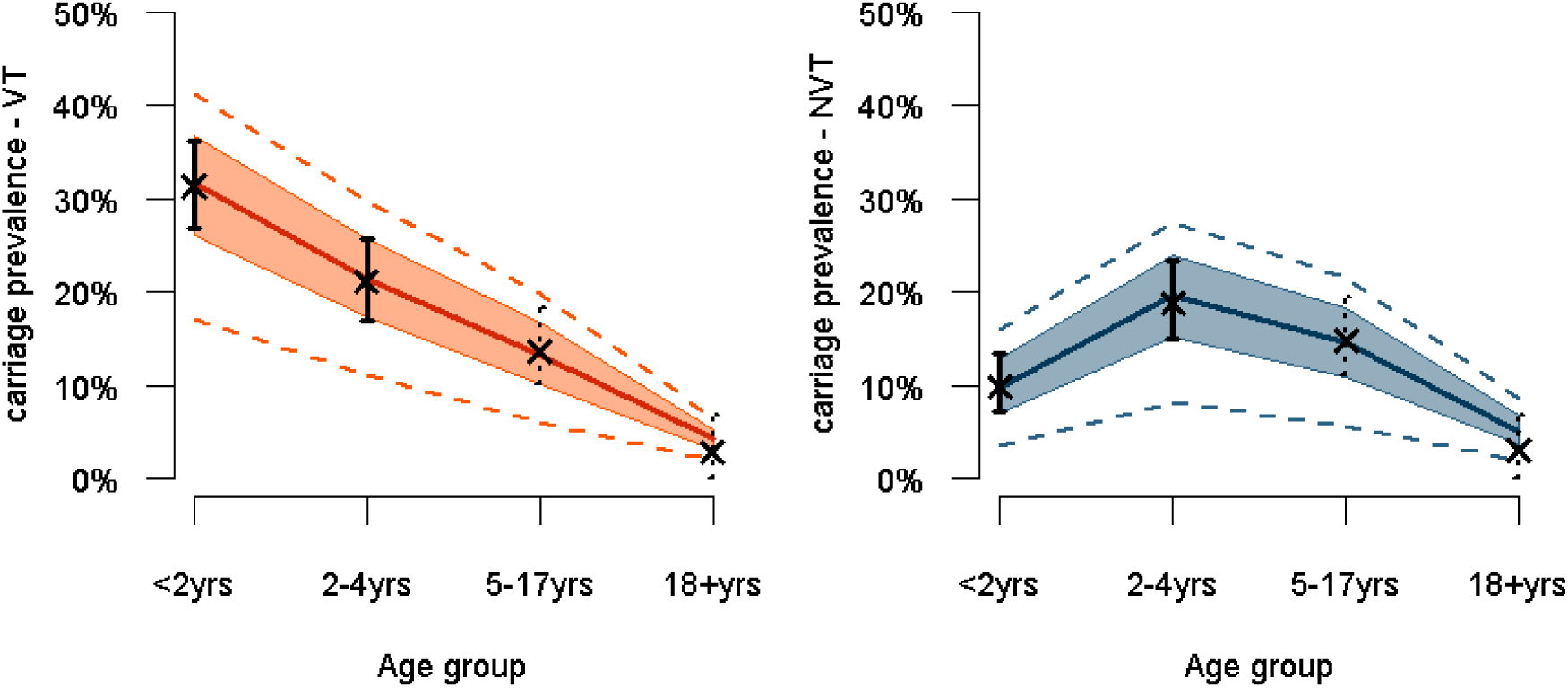
Pre-PCV carriage estimates across age groups for VT (left panel) and NVT (right panel), based on survey (plain vertical lines) and meta-regression model (dotted vertical lines) estimates, and estimates from the corresponding transmission model. **Legend:** Dots and bars correspond to the point and 95% confidence interval for the carriage estimates from the survey data (plain dot and plain lines) and the meta-regression model (cross and dotted lines). The dark red and dark blue plain lines represent the median transmission model estimate for VT and NVT respectively, the shaded areas the 50% credible interval (CrI) around the median and the dotted red and blue lines the 95%CrI

### Nasopharyngeal carriage

Our model predicts elimination of VT serotypes across all age groups within 10 years of PCV introduction in RV, with near-complete replacement by NVT serotypes, resulting in little or no change in the overall carriage prevalence (Figure 2A&B), particularly in children >5 years and in adults. CCs are predicted to reduce VT carriage more quickly through combined direct and indirect (i.e. herd) effects (Figures 2C and D). In <5 year olds the VT carriage in children is predicted to decrease by >99% within 6 years and 7 months (95% CI 5y2m − 9 y6m) in RV, 5 years and 11 months (4y6m– 8 y10m) in CC1, 5 years and 1 month (3y9m – 7y10m) in CC2 and 3 yrs (2y2m–4y10m) in CC5. Similar trends are predicted in older children and adults (Figures 2C and 2D).

**Figure 2:**
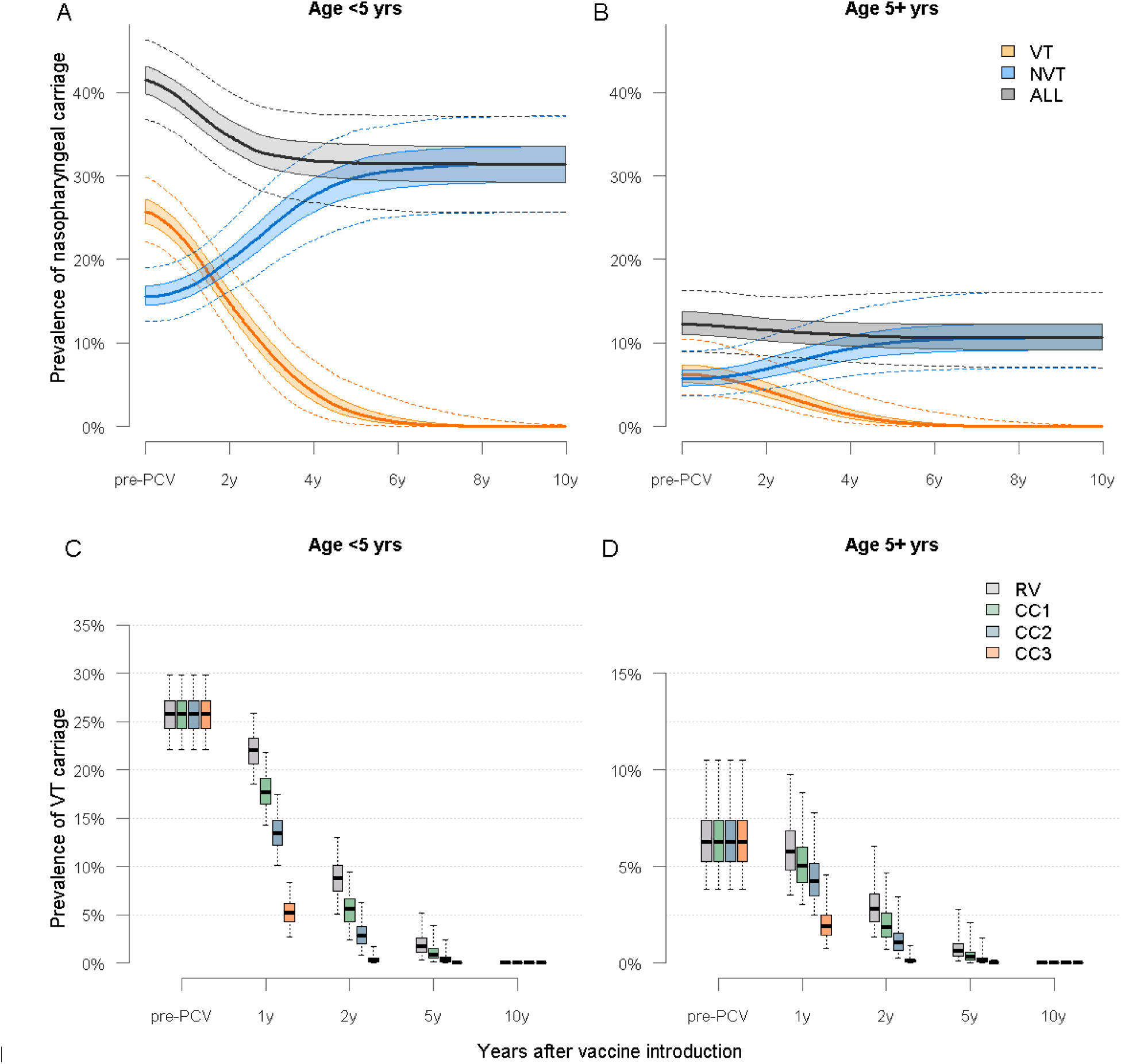
Predicted trends in nasopharyngeal carriage following PCV13 introduction in Nha Trang. **Legend:** <u>A</u>: Predicted trends in VT, NVT and overall carriage in <5 year olds without a catch-up campaign. <u>B</u>: Predicted trends in VT, NVT, and overall carriage in >5 year olds without a catch-up campaign. <u>C</u>: Predicted prevalence of VT carriage in <5 year olds for each vaccination strategy. <u>D</u>: Predicted prevalence of VT carriage in ≥5 year olds for each vaccination strategy. <u>In all four panels</u>: plain line= median, shaded areas= 50% credible intervals and dotted line or whiskers= 95% credible interval

### Invasive Pneumococcal Disease (IPD)

Our model predicts that the decline of IPD incidence will be proportionally highest among <2 year olds, falling to about 45% (95%CrI 33% − 57%) of its pre-PCV level and would be almost halved (57% (48 – 66%)) in children aged 2 – 4 years respectively (Figure 3A and 3B).

**Figure 3:**
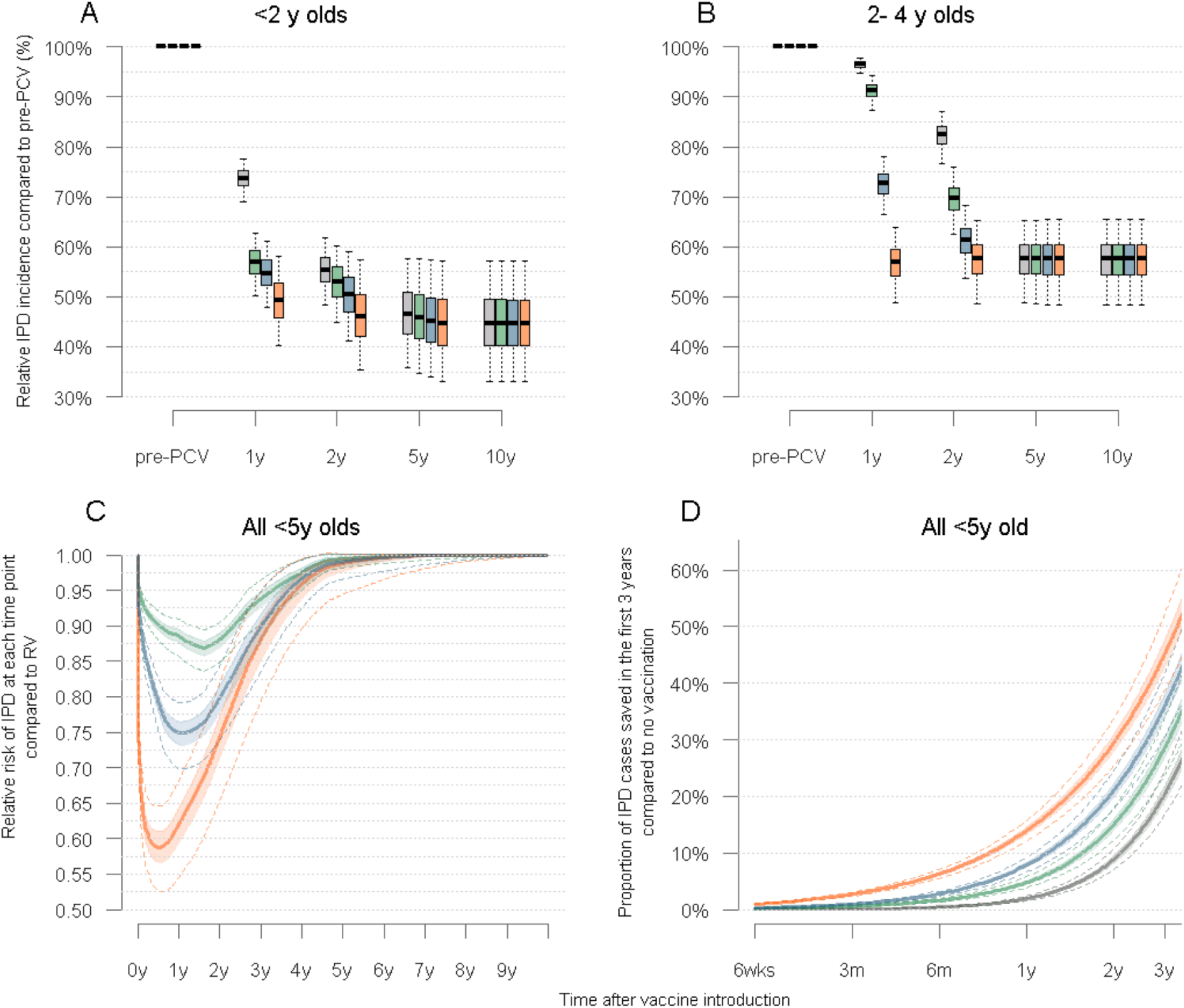
Trends in IPD following PCV introduction in children under five years of age in Nha Trang **Legend:** <u>A</u>: Predicted trends the cumulative annual incidence of IPD in children <2 years for each vaccination strategy considered, at a 90% vaccination coverage <u>B</u>: Predicted trends the cumulative annual incidence of IPD in children aged 2-4 years for each vaccination strategy considered, at a 90% vaccination coverage <u>C</u>: Cumulative number of IPD cases saved for each catch up strategy compared to RV, in children <5 years of age. <u>D</u>: Overall cumulative number of cases saved for each vaccination strategy, compared to no vaccination, in the first 3 years post PCV introduction. In all four panels: plain line= median, shaded areas= 50% credible intervals and dotted line or whiskers= 95% credible interval

Most of the benefit of CCs over routine vaccination in children <5 years is predicted to occur within the first three years after PCV introduction, with no noticeable difference at 5 years (Fig 3C). Our model predicts that the relative risk of IPD after CCs compared to RV would be lowest about 1 year after PCV introduction, with differences between strategies reducing thereafter until the new equilibrium is reached. Compared to RV, one year after vaccine introduction the number of cases of IPD is predicted to be 11% (95%CrI 9 – 14%) lower with CC1, 25% (21-30%) lower with CC2 and 38% (32 – 46%) lower with CC5 (Fig 3C).

The impact of each strategy on the cumulative proportion of cases averted in the first 3 years post PCV introduction, compared to no vaccination, is illustrated in Figure 3D.

Based on an average annual incidence risk of 49/100,000 children <5 years before PCV introduction [27], a routine introduction of PCV would result in a total of 74 cases (95%CrI − 86) per 100,000 children <5 years averted over the first five years of programme implementation and catch-up campaigns would lead to the prevention of an additional 13 (95%CrI 11 – 16) cases with CC1, 25 (95%CrI 21 – 30) cases with CC2 and 39 (95%CrI 31 – 49) cases with RV5.

### Delayed PCV introduction

We estimated the relative impact on IPD of catch-up campaigns for increasing delays. Our results suggest that, compared to RV, more IPD cases would be prevented in children <5 years insofar as PCV introduction is not delayed by more than 31 weeks (95%CI 30 – 32 weeks) for CC1, 58 weeks (53 – 63 weeks) for CC2 and 89 weeks (78 – 101 weeks) for CC5.

Vaccination delays would negatively impact <2 year olds more rapidly than 2 – 4 year old (Figure 4).

**Figure 4:**
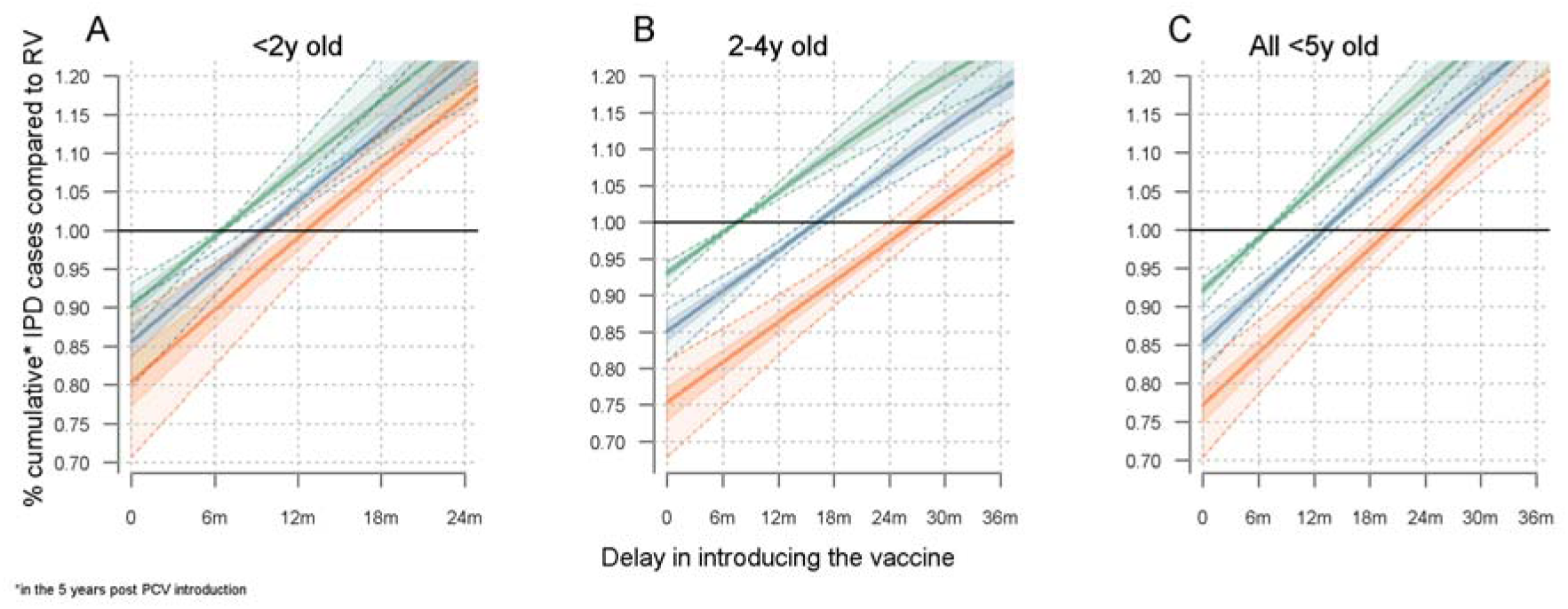
Impact of delayed PCV introduction with a catch-up campaign on VT carriage (panel A) and on IPD cases saved (panel B) in children under five years of age. **Legend:** <u>A</u>: <2 year olds. <u>B</u>: 2 − 4 year olds <u>C</u>: <5 years olds. The middle plain trend line corresponds to the median estimate (Green=CC1, Blue=CC2, Red=CC5), the dark shaded areas the 50% CrI and the light shaded areas the 95%CrI. The plain horizontal line at 1.00 represents the point below which interventions will be more favourable than a timely RV.

### Sensitivity analyses

#### Vaccination coverage

Our model predicts a lengthening of the time to near-elimination of VT serotypes (and hence, the time to reach the new post-PCV disease equilibrium) as vaccination coverage decreases but a similar differential impact of catch-up campaigns compared to routine vaccination. Full details are provided in the Supplementary File S3.

#### Duration of protection

A duration of protection of 3 years would increase the time to elimination of VT carriage, and thus prevent fewer IPD cases overall, while any average duration of protection longer than 6 years would not change model outcomes. With a duration of 3 years the median prevalence of VT in <5 year olds is predicted to reach near elimination about 2 years later in RV and CC1, 1.5 years later with CC2 and about 1 year later with CC5. Similar differences were predicted for the ≥5 year olds (Figure S5 in the Supplementary File S3). The relative impact of one vaccination strategy over another was predicted to be similar than with a duration of 6 years.

## DISCUSSION

We explored the possible impact of introducing PCV13 with and without a catch-up campaign in Vietnam through a dynamic transmission model. Our results feed into current debates about introduction strategies, particularly in South-East Asia where pneumococcal disease burden is high [28], where many countries have not yet introduced PCV [5], and where epidemiological data to guide decision making remain scarce [28, 29]. Although Vietnam is Gavi-eligible and is expected to introduce PCV in the coming years, it has not yet applied [5]. Our results provide estimates about how much catch-up campaigns would decrease disease burden compared to routine vaccine introduction without a campaign, for different scenarios. Our study also shows that, although catch-up campaigns would decrease disease burden more rapidly across age groups, their impact would only be beneficial insofar as the additional supply and operational constraints of their implementation does not delay PCV introduction by more than about 6 months to 2 years, depending on the age cohorts targeted by those campaigns.

The availability of data on both social mixing patterns – which are central to transmission models [13, 30] – and carriage in the same population allowed for thorough parameterisation of a transmission model.

However, our study has a number of limitations. In the absence of post-PCV data, predictions were based on the best available estimates of parameters governing vaccine effects that were observed elsewhere [16, 17, 31], which may not fully capture the local characteristics. Given that serotypes differ in their pathogenicity, fitness and transmissibility [32, 33], and that vaccine efficacy and duration of protection differs by serotype [12], our predictions based on homogeneous characteristics for the group of VT and NVT serotypes may overlook local epidemiological characteristics. This uncertainty was nonetheless captured to some extent by sampling from the known uncertainty around those parameters [16]. Moreover, we were not able to assess the impact of PCV on pneumococcal pneumonia, the burden of which is much higher than that of IPD [34], given the lack of robust data and the challenges in the aetiological assessment of clinical pneumonia. Results from ongoing studies in Gavi-eligible countries [35, 36] might help modelling work on the impact of PCV on pneumonia in the future.

Our predictions are in line with the experience of PCV7 in Europe and North America [37-44], as well with post-PCV trends observed in the few studies from low-income settings [36]. In the region of Kilifi in Kenya, where a catch-up campaign was conducted at PCV introduction, impact studies have shown a two third reduction in VT carriage prevalence across all age groups within two years of PCV10 introduction with a catch-up campaign among children <5 years [43].

The implementation of PCV in various settings has consistently resulted in little or no change in overall carriage prevalence, due to replacement effects by NVT serotypes colonising the space left vacant by VT in the nasopharynx, but a reduction in severe disease given the lower pathogenicity of the latter, in accordance with our model output. Our predictions were also robust to estimates of duration of protection of vaccination coverage, and thus provide useful estimates of the impact of introducing PCV in a semi-urban Southeast Asian setting.

While supply issues could also lead to other problems than just delay, such as incomplete schedules once a programme has started, we here assumed that the planning of vaccine introduction would be based on vaccine availability, full roll-out capacity and sustainability at introduction. With supply constraints, and ignoring other supply side, staff and outreach challenges that could potentially delay the implementation of CCs [5], our study can inform whether delaying the introduction of the vaccine to allow for CCs would potentially be beneficial, compared to a routine-only strategy. Moreover, this model also provides a framework that could feed into economic evaluations to further guide decisions about vaccine introduction with and without campaigns.

The generalisability to other settings of the differential impact of CCs needs to be considered in light of the epidemiological and socio-demographic characteristics of Nha Trang. In particular, the low prevalence of carriage and an ageing population - with only 5% of children under the age of five years – impact on the speed with which herd effects are established. Although similar epidemiological and demographic characteristics are observed in many other South(east) Asian settings [45, 46], the differential impact of CCs and the establishment of herd effects in settings with a younger population and a higher carriage prevalence is likely to differ, and should be addressed with models applied to such settings.

In conclusion, our study offers insights into the current debate about vaccination strategies when introducing PCV in South-East Asia. Our model suggests that catch-up campaigns have the potential to rapidly decrease carriage and disease across age groups, but are only offering added reduction in disease burden insofar their implementation results in little to no implementation delay.

## DECLARATIONS

### Ethics approval

Ethical approval for the field surveys was granted by the institutional review board of the London School of Hygiene and Tropical Medicine and the Ethics Commission of National Institute of Hygiene and Epidemiology, Hanoi and the Nagasaki University.

Consent for publication

### Availability of data and material

Other model outputs are available on request. Please contact the corresponding author at

### Authors’ contributions

#### Acknowledgements

We would like to thank Dr Ana Maria Henao Restrepo, Dr Hope Johnson and Prof Katherine L O’Brien for helpful discussions around the study objectives.

We are grateful to the participants of this study and their parents as well as to the staff from Khanh-Hoa Health Service and the medical staff from Khanh Hoa General Hospital for their support. We also thank the staff from the Japan-Vietnam Friendship Laboratory at National Institute of Hygiene and Epidemiology, Hanoi, and Institute of Tropical Medicine, Nagasaki University.

Authors’ information (optional)

## LIST OF ABBREVIATIONS

CC: Catch-up campaign
CC1: CC in <1 year olds
CC2: CC in <2 year olds
CC5: CC in <5 year olds
IPD: Invasive Pneumococcal Disease
MCMC: Markov Chain Monte-Carlo
NVT: Non vaccine type
PCV: pneumococcal conjugate vaccine
PCV7: seven-valent PCV
PCV10: 10-valent PCV
PCV13: 13-valent PCV
RV: Routine vaccination
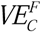: vaccine efficacy against carriage conferring full protection
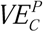: vaccine efficacy against carriage conferring partial protection
VT: Vaccine Type

## Supplementary File S1:Model structure

We built a realistic age-structured deterministic Susceptible-Infected-Susceptible (SIS) transmission model of carriage acquisition and clearance, similar to Choi et al. [16].

The model considers all VT serotypes jointly and separately from the group of NVT serotypes, but allows for VT-NVT co-colonization, thus resulting in four compartments of carriage states, namely susceptible, VT carriers, NVT carriers and VT-NVT co-colonized as in Choi et al. [16]. In this model movements from the susceptible to the VT or NVT compartments are determined by age-specific forces of infection for VT (λ_Vi_) and NVT (λ_Ni_) respectively, and co-colonization is determined by competition parameters (C_N_ and C_V_), which represent the degree with which prior colonization reduces the likelihood of co-colonization. Age-specific recovery rates (r_i_) determine the speed with which individuals revert back from the co-colonized to either VT or NVT compartments, and from colonization to susceptible compartments, assuming no natural immunity to carriage.

The model comprised of three levels of vaccine-induced immunity; (1) no protection, (2) partial protection and (3) full protection (Figure S1). The latter means the efficacy and duration of protection conferred after completion of the infant schedule (i.e. 2 infant doses and a booster at 12 months (‘2+1’ schedule)) or the completion of a catch-up programme in older children (2 doses in <18 months and 1 dose in ≥18 months). Partial protection was gained form two primary infant doses, or after the first catch-up dose in children aged 12 - 17 months. The difference between full and partial protection lied in the magnitude of vaccine efficacy against carriage (*VE_C_*) and in the duration of protection.

We applied the model to a population of 81 annual age cohorts (0 to 80 years) divided into weekly age bands of 100 individuals. In the calculation of the force of infection the population figure was adjusted to represent the actual population, based on census data.

We inferred the impact on IPD based posterior estimates of carriage at each time step, case:carrier ratios for VT and NVT by age group [16], the vaccine efficacy against invasiveness and the vaccination coverage of each weekly age cohort at each time step. Details about parameters of vaccine efficacy are provided in the Supplementary File S2.

The model equations are provided in Text S1 below.

**Figure S1:**
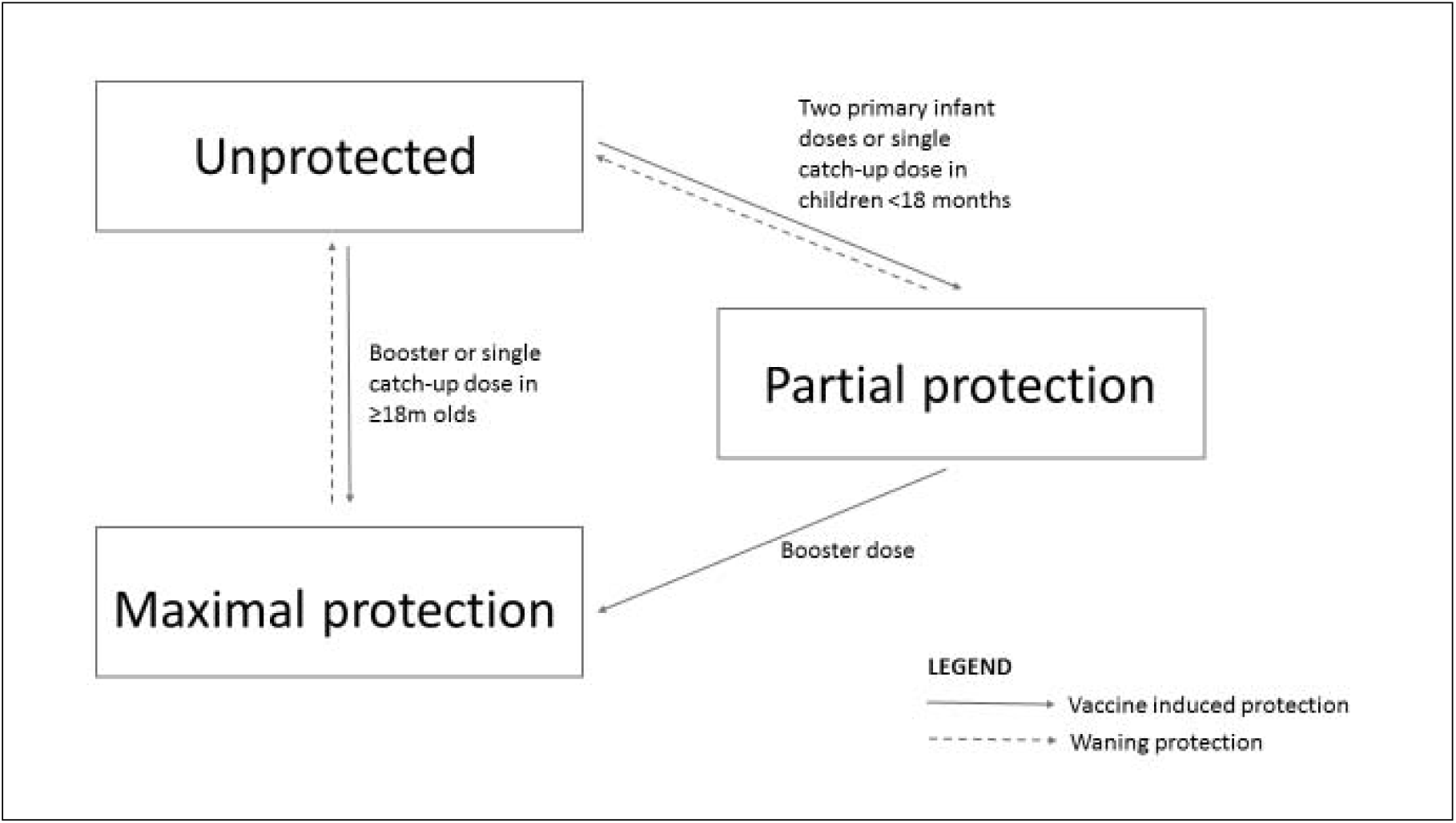
The three groups of vaccine protection states defined in the model and movement between groups

**Text S1**: Model equations

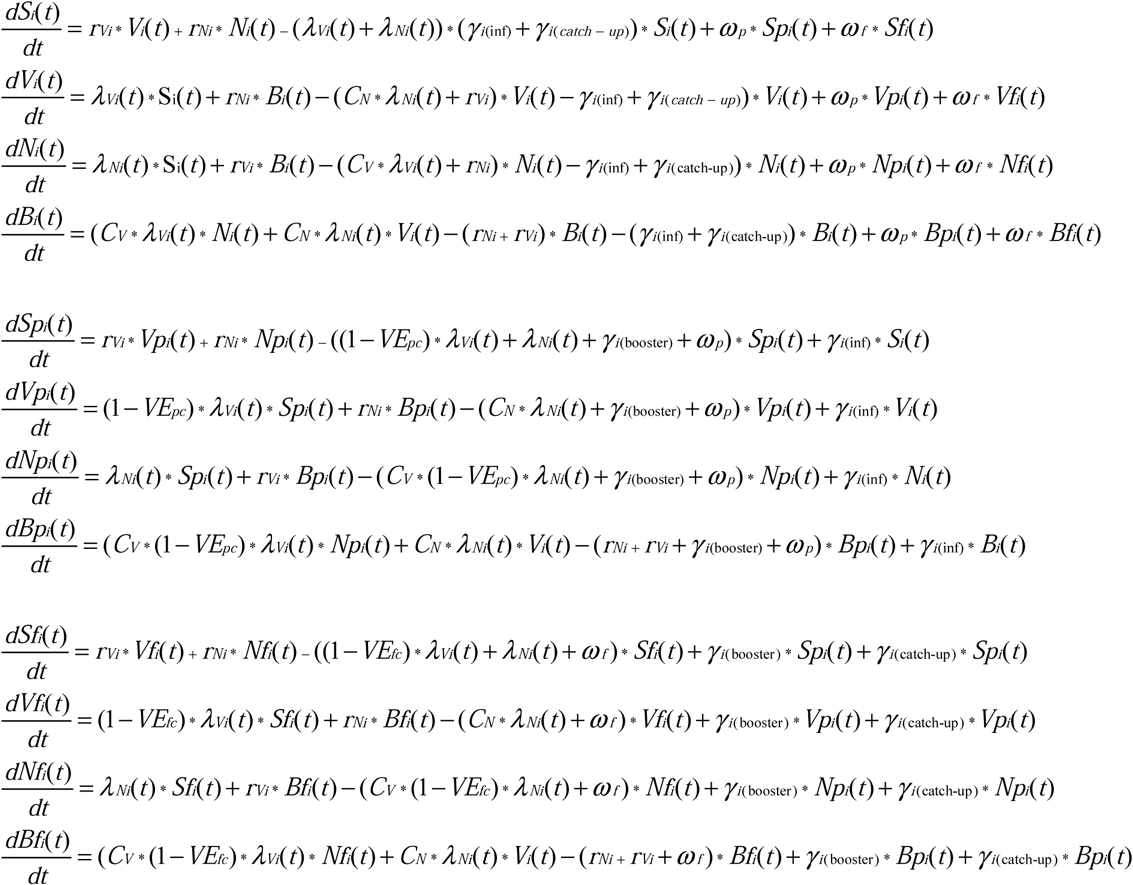

*i* represent the weekly age groups. is the vaccination coverage for infant doses (inf), the booster dose (boost) and the catch-up dose (catch-up). VE*f*_C_ and VE_*p*C_ is the vaccine efficacy against carriage acquisition after full and partial vaccination respectively. ω_P_ and ω_f_ represent the rate of waning immunity after partial and full vaccination.

## Supplementary File S2:Vaccine efficacy and duration of protection

### Vaccine efficacy against carriage

#### Full vaccination

We obtained the vaccine efficacy against carriage after full protection (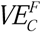) and its uncertainty from a Bayesian meta-regression model based on estimates from 22 intervention studies. The details have been published elsewhere [17]. The model of vaccine efficacy was run for as many iterations as in our transmission model, so that we could sample one estimate of 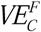 at each iteration of the post-vaccination simulations.

#### Partial vaccination

A systematic review of the impact of pneumococcal conjugate vaccines on carriage [50] provided information on intervention studies with 2 primary-dose arms where compared with either 2+1 schedules or 3+0 schedules.

We included individual randomized controlled trials providing nasopharyngeal carriage estimates within 3 – 20 months after a 2-dose infant schedule (2+0 arm) and after a complete schedule (either 3+0 or 2+1 arm). We excluded studies providing carriage estimates within 3 months after vaccination to account for the delay in producing an immune response, and time to carriage clearance under the assumption that existing carriage at the time of vaccination is not affected by PCV [17].

We extracted data on the number of carriers of VT and total number of individuals swabbed in each study arm before and after vaccination. We then computed the relative risk of VT carriage among fully vaccinated compared to partially vaccinated children. The pooled estimates were obtained through a random effects meta-regression model.

We pooled estimates from four studies, including trials in Fiji [19], Israel[18], The Gambia [51] and The Netherlands [20], providing together eight different survey points, into a simple random effects meta-analysis, not taking into account within-study dependence given the limited number of data points per study. The details of the studies are provided in Table S1 below.

**Table S1:** Studies included in the meta-analysis.

| Study | Country | Schedule | Age<br>(months) | Time since<br>last dose<br>(months) | Carriers /total | Relative Risk (95%) |
| --- | --- | --- | --- | --- | --- | --- |
| Dagan (2012)[18]* | Israel | 3+0 | 9.5m | 3.5m | 58/327 | 0.83 (0.57 – 1.20) |
| Dagan (2012)[18]* | Israel | 2+0 | 9.5m | 3.5m | 36/169 | ref |
| Ota (2011)[51] | Gambia | 3+0 | 11m | 7m | 20/200 | 0.59 (0.36 – 1.00) |
| Ota (2011)[51] | Gambia | 2+0 | 11m | 8m | 33/198 | ref |
| Ota (2011)[51] | Gambia | 3+0 | 15m | 11m | 24/193 | 0.80 (0.49 – 1.32) |
| Ota (2011)[51] | Gambia | 2+0 | 15m | 12m | 30/196 | ref |
| Russell (2010)[19] | Fiji | 3+0 | 6m | 3m | 13/127 | 0.95 (0.47 – 1.89) |
| Russell (2010)[19] | Fiji | 2+0 | 6m | 3m | 16/148 | ref |
| Russell (2010)[19] | Fiji | 3+0 | 9m | 6m | 4/122 | 0.32 (0.11 – 0.93) |
| Russell (2010)[19] | Fiji | 2+0 | 9m | 6m | 15/146 | ref |
| Russell (2010)[19] | Fiji | 3+0 | 12m | 9m | 8/114 | 1.25 (0.49 – 3.23) |
| Russell (2010)[19] | Fiji | 2+0 | 12m | 9m | 9/143 | ref |
| van Gils (2009)[20] | Netherlands | 2+1 | 18m | 7m | 51/329 | 0.64 (0.47 – 0.88) |
| van Gils (2009)[20] | Netherlands | 2+0 | 18m | 14m | 79/327 | ref |
| van Gils (2009)[20] | Netherlands | 2+1 | 24m | 13m | 47/333 | 0.96 (0.66 -1.39) |
| van Gils (2009)[20] | Netherlands | 2+0 | 24m | 20m | 49/332 | ref |

\*In this study pooled results were provided for samples taken at 7 months and 12 months of age, and we therefore considered the age and time since the last dose as the average between the two

The pooled relative risk of carriage among children with a full schedule was 78% (95%CI 64 – 92%) that of a partial schedule (Figure S2), with no evidence of heterogeneity (I^2^=3%).

At each iteration of the MCMC, we sampled a value from the relative risk 78% (95%CI 64 – 92%) multiplied by a value of the full vaccine efficacy against carriage acquisition (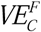), to obtain a measure of the partial efficacy (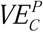).

**Figure S2:**
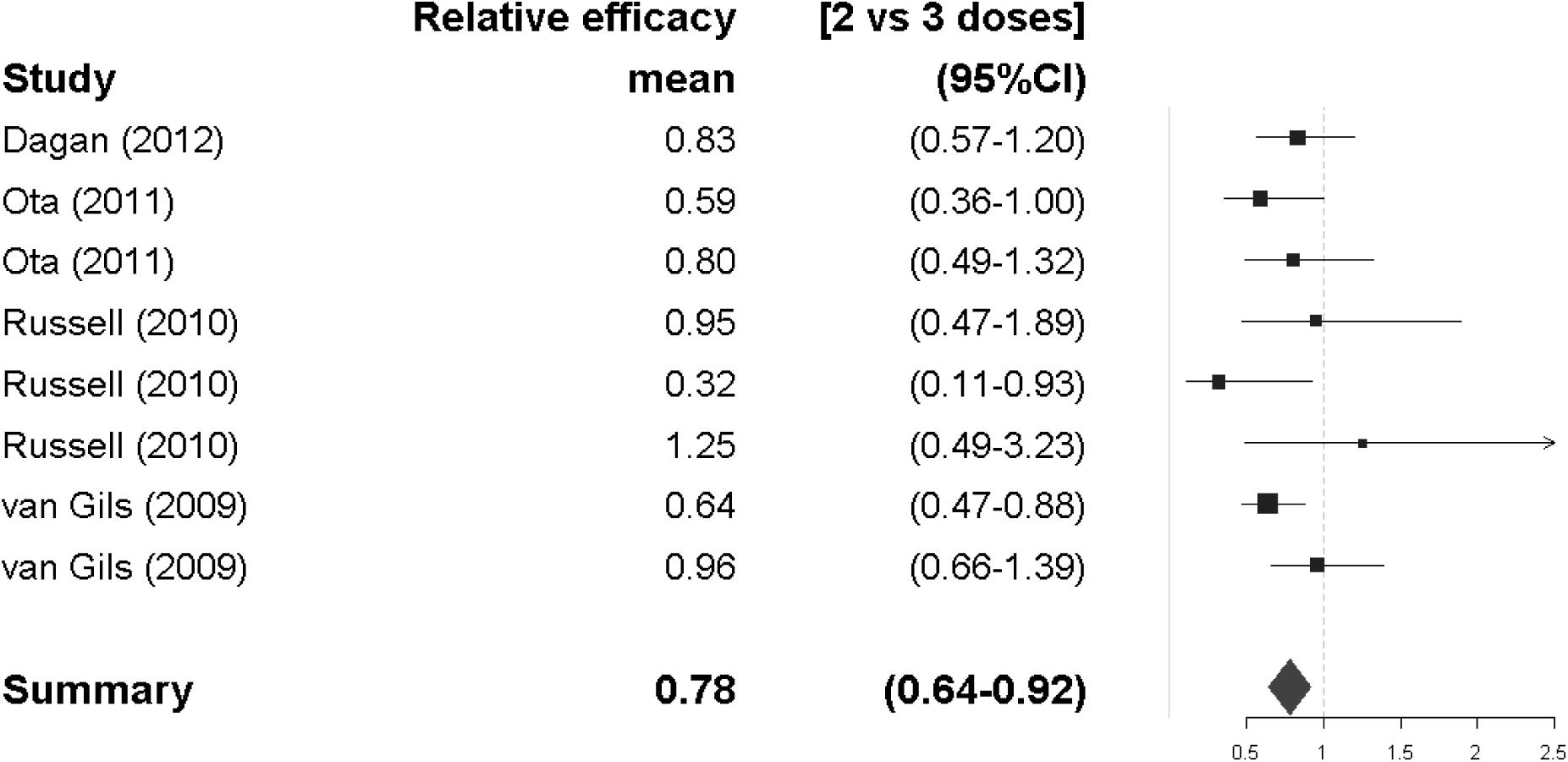
Forest plot of the relative risk of VT carriage of 3 primary doses (2+1 or 3+0), for four different individual randomized controlled trials (RCTs), at eight different time points [18-20, 51].

### Duration of protection

The duration of protection against carriage acquisition was obtained from le Polain et al [17]. The median estimate of a model of waning efficacy, using an asymptomatic function, closely matched that of an exponential decay with a mean duration of protection of 6 years (Figure S3). Hence, this estimate was taken as the average duration of protection in the main model. For sensitivity analyses, we considered mean durations of 3 years and 20 years as the lower and higher values, as shown in Figure S3.

**Figure S3:**
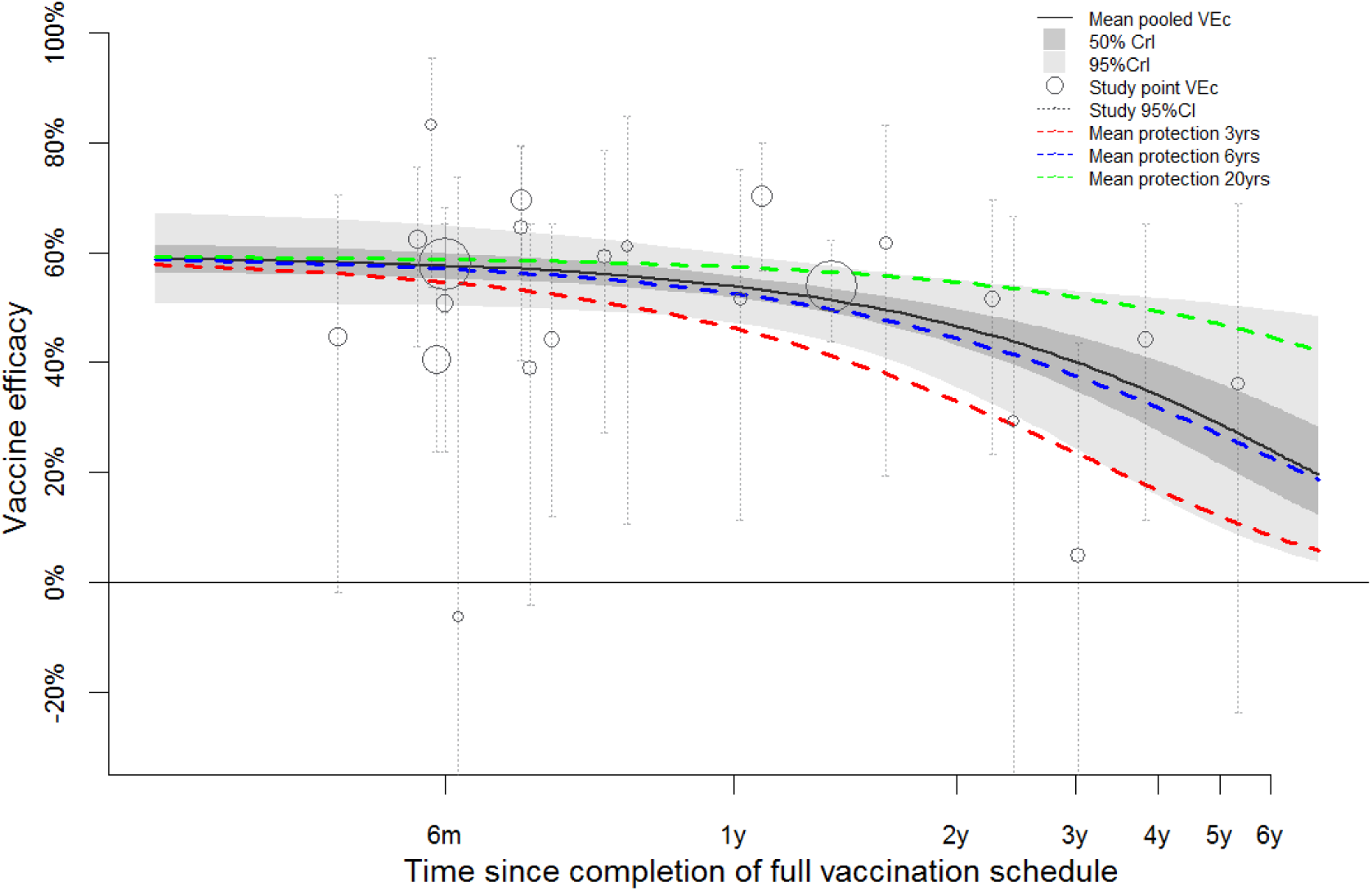
Duration of protection of PCV against carriage acquisition

**Legend**: Each circle represents the mean vaccine efficacy estimate for each study included in the analysis – see [17]. The size of the circle is proportional to the study size. The whiskers on either side of each circle represent the 95%CrI around the point estimate. The dark grey area corresponds to the 50% CrI around the main model estimate of vaccine efficacy and its waning (plain black line), and the lighter grey area to the 95% CrI. The blue, red and green dotted lines show the waning with an exponential decay function for a mean duration of protection of 6 years, 3 years and 20 years respectively. Figure adapted from le Polain de Waroux et al. [17], with permission from PIDJ

In the absence of estimates on the duration of protection of partial vaccination, we assumed that the duration of protection of a partial vaccination was 0.78 (0.64 – 0.92) that of a full vaccination, as for VE_C_.

### Vaccine efficacy against Invasive Pneumococcal Disease

The vaccine efficacy against invasive pneumococcal disease (VE_IPD_) can be expressed as a function of the vaccine efficacy against carriage acquisition (VE_C_) and the efficacy against progression to disease as a result of carriage, which we here term the vaccine efficacy against invasiveness (VE_inv_) [25], where VE_IPD_ = 1 − (1-VE_C_)(1-VE_inv_).

Based on estimates from a large systematic review by Lucero et al. [26], we assumed VE_IPD_ to be 80% (95%CI 58 – 90%) and generated a binomial distribution that closely matched those estimates. Hence, we calculated VE_inv_ based on the distribution of VE_IPD_ and that of VE_C_, and obtained a value of VE_inv_ at each iteration in our MCMC process.

The mean VE_INV_ was 80% (95%CrI 67 – 90%), the mean VE_C_ was 62% (95%CrI 52 – 73%) and mean VE_inv_ was 48% (95%CrI 34 – 65%). The distributions are shown in Figure S4 below.

**Figure S4:**
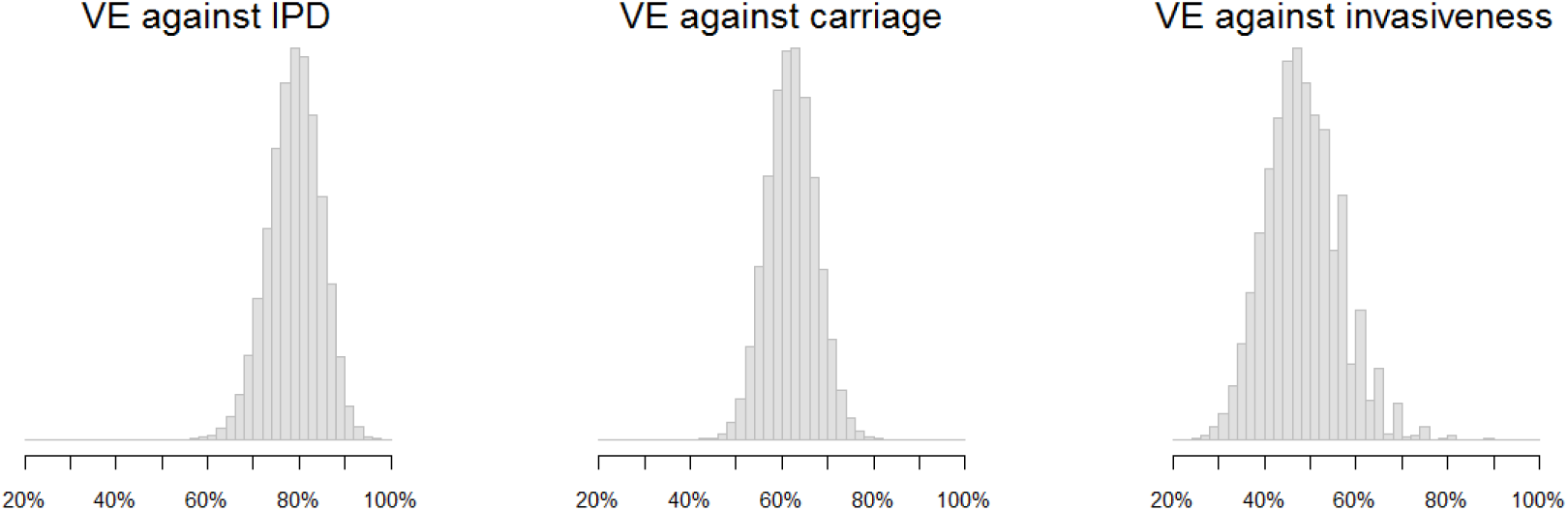
Estimates of vaccine efficacy

**Legend**: density plot of the vaccine efficacy against IPD (left panel), carriage (medium panel) and invasiveness (right panel). The x axis displays the vaccine efficacy.

In the model, was assumed no waning of VE_inv_, and that VE_inv_ would be conferred after 2 primary doses, or any catch-up dose. Although there is evidence that VE_C_ after partial vaccination differs from that after complete vaccination (see earlier), evidence that the efficacy against progression to invasiveness differs between two and three primary doses is scarce.

Although it is likely that VE_inv_ wanes over time, we did not consider waning in the main model, given that most of the direct impact of PCV on IPD occurs in the first few years of life, before establishment of the herd immunity effect (see results), and given the lack of estimates of waning of VE_INV_.

## Supplementary File S3: Sensitivity analysis (coverage and duration of vaccine protection)

### Vaccination coverage

We explored model outputs with lower vaccination coverage in both cohort and catch-up immunization. Our model predicted a lengthening of the time to near-elimination of VT serotypes (and hence, the time reach the new post-PCV disease equilibrium) as vaccination coverage lowers but a similar differential impact of catch-up campaigns compared to routine vaccination.

Figure S4 compares predictions of trends in VT carriage for all four strategies and vaccination coverage levels of 90%, 70% and 50%.

### Duration of protection

A duration of protection of 3 years would increase the time to elimination of VT carriage, and thus prevent fewer IPD cases overall, while any average duration of protection longer than 6 years would not change model outcomes. With a duration of 3 years the median prevalence of VT in <5 year olds is predicted to reach near elimination about 2 years later in RV and CC1, 1.5 years later with CC2 and about 1 year later with CC5. Similar differences were predicted for the ≥5 year olds (Figure S5). The relative impact of one vaccination strategy over another was predicted to be similar than with a duration of 6 years.

**Figure S4:**
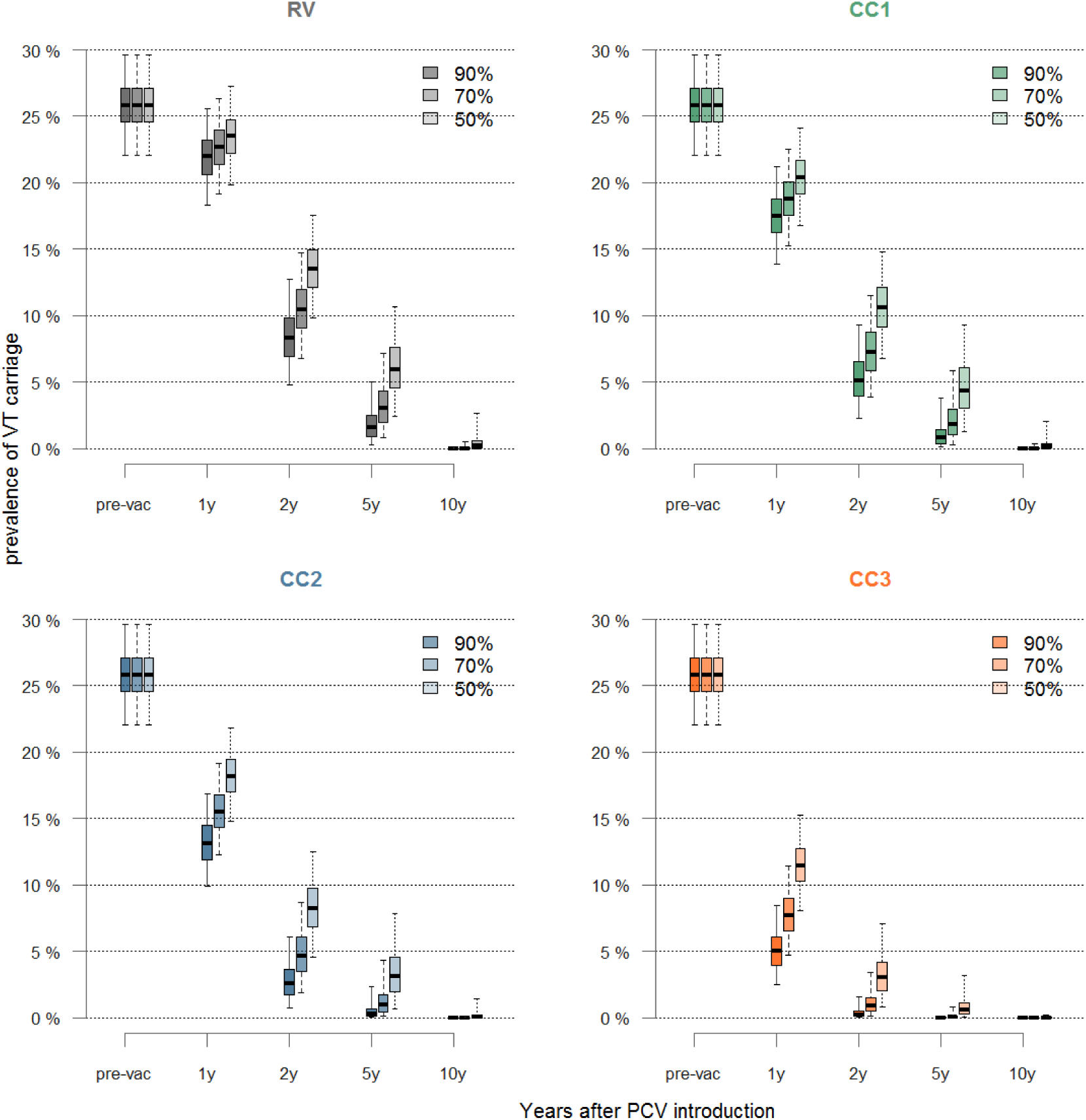
Impact of vaccination coverage on VT carriage for each strategy considered, and comparing 90%, 70% and 50% homogeneous coverage for each of the four strategies, in children aged 0 – 59 months

<u>Legend</u>: In all four panels: plain horizontal line= median, boxes= 50% credible intervals and whiskers= 95% credible interval, for each of the three levels of vaccination coverage (90%,70%, 50%) and for all four vaccination strategies.

**Figure S5:**
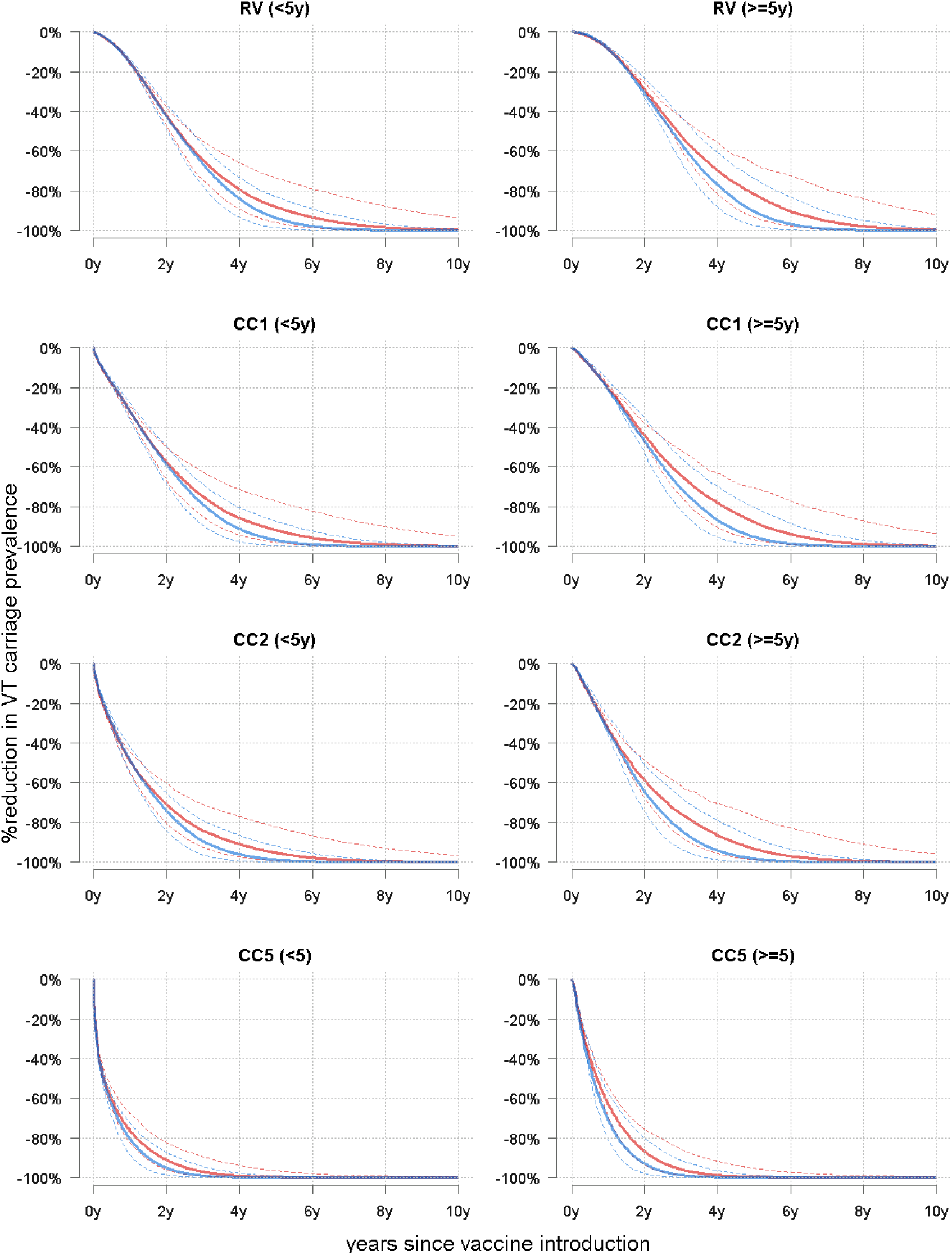
Comparing model estimates with an average 905 duration of protection of 6 years (blue) and of 3 years (red) for each of the vaccination scenarios considered and by age group

<u>Legend:</u> Percentage reduction in VT carriage in children <5 years (left) and ≥ 5 years old. In blue is are estimates of the main model (average duration of protection of 6 years) and in red are estimates considering a mean duration of protection of 3 years. The plain lines represent the median and the dotted lines the 95% credible interval

## REFERENCES

[1] Cohen AL, Hyde TB, Verani J, Watkins M. Integrating pneumonia prevention and treatment interventions with immunization services in resource-poor countries. Bulletin of the World Health Organization. 2012;90:289–94.

[2] Black RE, Cousens S, Johnson HL, Lawn JE, Rudan I, Bassani DG, et al. Global, regional, and national causes of child mortality in 2008: a systematic analysis. Lancet. 2010;375:1969–87.

[3] Global Action Plan for prevention and control of pneumonia (GAPP). Geneva: World Health Organization; 2008.

[4] Calix JJ, Porambo RJ, Brady AM, Larson TR, Yother J, Abeygunwardana C, et al. Biochemical, genetic, and serological characterization of two capsule subtypes among Streptococcus pneumoniae Serotype 20 strains: discovery of a new pneumococcal serotype. J Biol Chem. 2012;287:27885–94.

[5] Gavi, The Vaccine Alliance,. Advance Market Commitment for Pneumococcal Vaccines. Annual Report 1 April 2014 - 31 March 2015. In: Gavi TVA, editor.2015.

[6] Harris JB, Gacic-Dobo M, Eggers R, Brown DW, Sodha SV, Centers for Disease C, et al. Global routine vaccination coverage, 2013. MMWR Morbidity and mortality weekly report. 2014;63:1055–8.

[7] Publication WHO. Pneumococcal vaccines WHO position paper - 2012 - recommendations. Vaccine. 2012;30:4717–8.

[8] Whitney CG, Goldblatt D, O’Brien KL. Dosing schedules for pneumococcal conjugate vaccine: considerations for policy makers. The Pediatric infectious disease journal. 2014;33 Suppl 2:S172–81.

[9] Vu HT, Yoshida LM, Suzuki M, Nguyen HA, Nguyen CD, Nguyen AT, et al. Association between nasopharyngeal load of Streptococcus pneumoniae, viral coinfection, and radiologically confirmed pneumonia in Vietnamese children. The Pediatric infectious disease journal. 2011;30:11–8.

[10] Yoshida LM, Nguyen HA, Watanabe K, Le MN, Nguyen AT, Vu HT, et al. Incidence of radiologically-confirmed pneumonia and Haemophilus influenzae type b carriage before Haemophilus influenzae type b conjugate vaccine introduction in Central Vietnam. The Journal of pediatrics. 2013;163:S38–43.

[11] O’Brien KL, Nohynek H, World Health Organization Pneumococcal Vaccine Trials Carriage Working G. Report from a WHO Working Group: standard method for detecting upper respiratory carriage of Streptococcus pneumoniae. The Pediatric infectious disease journal. 2003;22:e1–11.

[12] Le Polain de Waroux O, Flasche S, Prieto-Merino D, Edmunds WJ. Age-Dependent Prevalence of Nasopharyngeal Carriage of Streptococcus pneumoniae before Conjugate Vaccine Introduction: A Prediction Model Based on a Meta-Analysis. PloS one. 2014;9:e86136.

[13] Mossong J, Hens N, Jit M, Beutels P, Auranen K, Mikolajczyk R, et al. Social contacts and mixing patterns relevant to the spread of infectious diseases. PLoSMed. 2008;5:e74.

[14] Melegaro A, Jit M, Gay N, Zagheni E, Edmunds WJ. What types of contacts are important for the spread of infections?: using contact survey data to explore European mixing patterns. Epidemics. 2011;3:143–51.

[15] Kadioglu A, Weiser JN, Paton JC, Andrew PW. The role of Streptococcus pneumoniae virulence factors in host respiratory colonization and disease. Nature reviews Microbiology. 2008;6:288–301.

[16] Choi YH, Jit M, Gay N, Andrews N, Waight PA, Melegaro A, et al. 7-Valent pneumococcal conjugate vaccination in England and Wales: is it still beneficial despite high levels of serotype replacement? PLoSOne. 2011;6:e26190.

[17] Le Polain De Waroux O, Flasche S, Prieto-Merino D, Goldblatt D, Edmunds WJ. The Efficacy and Duration of Protection of Pneumococcal Conjugate Vaccines Against Nasopharyngeal Carriage: A Meta-regression Model. The Pediatric infectious disease journal. 2015;34:858–64.

[18] Dagan R, Givon-Lavi N, Porat N, Greenberg D. The effect of an alternative reduced-dose infant schedule and a second year catch-up schedule with 7-valent pneumococcal conjugate vaccine on pneumococcal carriage: a randomized controlled trial. Vaccine. 2012;30:5132–40.

[19] Russell FM, Carapetis JR, Satzke C, Tikoduadua L, Waqatakirewa L, Chandra R, et al. Pneumococcal nasopharyngeal carriage following reduced doses of a 7-valent pneumococcal conjugate vaccine and a 23-valent pneumococcal polysaccharide vaccine booster. Clinical and vaccine immunology: CVI. 2010;17:1970–6.

[20] van Gils EJ, Veenhoven RH, Hak E, Rodenburg GD, Bogaert D, Ijzerman EP, et al. Effect of reduced-dose schedules with 7-valent pneumococcal conjugate vaccine on nasopharyngeal pneumococcal carriage in children: a randomized controlled trial. JAMA: the journal of the American Medical Association. 2009;302:159–67.

[21] Ferraro CF, Trotter CL, Nascimento MC, Jusot JF, Omotara BA, Hodgson A, et al. Household crowding, social mixing patterns and respiratory symptoms in seven countries of the African meningitis belt. PloS one. 2014;9:e101129.

[22] Melegaro A, Choi Y, Pebody R, Gay N. Pneumococcal carriage in United Kingdom families: estimating serotype-specific transmission parameters from longitudinal data. American journal of epidemiology. 2007;166:228–35.

[23] Ali M, Canh GD, Clemens JD, Park JK, von Seidlein L, Minh TT, et al. The use of a computerized database to monitor vaccine safety in Viet Nam. Bulletin of the World Health Organization. 2005;83:604–10.

[24] Choi YH, Jit M, Flasche S, Gay N, Miller E. Mathematical modelling long-term effects of replacing Prevnar7 with Prevnar13 on invasive pneumococcal diseases in England and Wales. PLoSOne. 2012;7:e39927.

[25] Simell B, Auranen K, Kayhty H, Goldblatt D, Dagan R, O’Brien KL. The fundamental link between pneumococcal carriage and disease. Expert review of vaccines. 2012;11:841–55.

[26] Lucero MG, Dulalia VE, Nillos LT, Williams G, Parreno RA, Nohynek H, et al. Pneumococcal conjugate vaccines for preventing vaccine-type invasive pneumococcal disease and X-ray defined pneumonia in children less than two years of age. Cochrane database of systematic reviews. 2009:CD004977.

[27] Anh DD, Kilgore PE, Slack MP, Nyambat B, Tho le H, Yoshida LM, et al. Surveillance of pneumococcal-associated disease among hospitalized children in Khanh Hoa Province, Vietnam. Clinical infectious diseases: an official publication of the Infectious Diseases Society of America. 2009;48 Suppl 2:S57–64.

[28] O’Brien KL, Wolfson LJ, Watt JP, Henkle E, Deloria-Knoll M, McCall N, et al. Burden of disease caused by Streptococcus pneumoniae in children younger than 5 years: global estimates. Lancet. 2009;374:893–902.

[29] Jaiswal N, Singh M, Das RR, Jindal I, Agarwal A, Thumburu KK, et al. Distribution of serotypes, vaccine coverage, and antimicrobial susceptibility pattern of Streptococcus pneumoniae in children living in SAARC countries: a systematic review. PloS one. 2014;9:e108617.

[30] Wallinga J, Teunis P, Kretzschmar M. Using data on social contacts to estimate age-specific transmission parameters for respiratory-spread infectious agents. AmJEpidemiol. 2006;164:936–44.

[31] Auranen K, Mehtala J, Tanskanen A, Kaltoft S. Between-strain competition in acquisition and clearance of pneumococcal carriage‐‐epidemiologic evidence from a longitudinal study of day-care children. AmJEpidemiol. 2010;171:169–76.

[32] Lipsitch M, Abdullahi O, D’Amour A, Xie W, Weinberger DM, Tchetgen Tchetgen E, et al. Estimating rates of carriage acquisition and clearance and competitive ability for pneumococcal serotypes in Kenya with a Markov transition model. Epidemiology. 2012;23:510–9.

[33] Hogberg L, Geli P, Ringberg H, Melander E, Lipsitch M, Ekdahl K. Age- and serogroup-related differences in observed durations of nasopharyngeal carriage of penicillin-resistant pneumococci. Journal of clinical microbiology. 2007;45:948–52.

[34] Rudan I, El Arifeen S, Bhutta ZA, Black RE, Brooks A, Chan KY, et al. Setting research priorities to reduce global mortality from childhood pneumonia by 2015. PLoS medicine. 2011;8:e1001099.

[35] Levine OS, Bhat N, Crawley J, Deloria-Knoll M, DeLuca AN, Driscoll AJ, et al. Pneumonia etiology research for child health. Introduction. Clinical infectious diseases: an official publication of the Infectious Diseases Society of America. 2012;54 Suppl 2:S87–8.

[36] Rodgers GL, Klugman KP. Surveillance of the impact of pneumococcal conjugate vaccines in developing countries. Hum Vaccin Immunother. 2016;12:417–20.

[37] Flasche S, van Hoek AJ, Sheasby E, Waight P, Andrews N, Sheppard C, et al. Effect of pneumococcal conjugate vaccination on serotype-specific carriage and invasive disease in England: a cross-sectional study. PLoSMed. 2011;8:e1001017.

[38] Huang SS, Hinrichsen VL, Stevenson AE, Rifas-Shiman SL, Kleinman K, Pelton SI, et al. Continued impact of pneumococcal conjugate vaccine on carriage in young children. Pediatrics. 2009;124:e1–11.

[39] Spijkerman J, van Gils EJ, Veenhoven RH, Hak E, Yzerman EP, van der Ende A, et al. Carriage of Streptococcus pneumoniae 3 years after start of vaccination program, the Netherlands. EmergInfectDis. 2011;17:584–91.

[40] Ghaffar F, Barton T, Lozano J, Muniz LS, Hicks P, Gan V, et al. Effect of the 7-valent pneumococcal conjugate vaccine on nasopharyngeal colonization by Streptococcus pneumoniae in the first 2 years of life. Clinical infectious diseases: an official publication of the Infectious Diseases Society of America. 2004;39:930–8.

[41] Pelton SI, Loughlin AM, Marchant CD. Seven valent pneumococcal conjugate vaccine immunization in two Boston communities: changes in serotypes and antimicrobial susceptibility among Streptococcus pneumoniae isolates. The Pediatric infectious disease journal. 2004;23:1015–22.

[42] Hennessy TW, Singleton RJ, Bulkow LR, Bruden DL, Hurlburt DA, Parks D, et al. Impact of heptavalent pneumococcal conjugate vaccine on invasive disease, antimicrobial resistance and colonization in Alaska Natives: progress towards elimination of a health disparity. Vaccine. 2005;23:5464–73.

[43] Hammitt LL, Bruden DL, Butler JC, Baggett HC, Hurlburt DA, Reasonover A, et al. Indirect effect of conjugate vaccine on adult carriage of Streptococcus pneumoniae: an explanation of trends in invasive pneumococcal disease. JInfectDis. 2006;193:1487–94.

[44] Flasche S, Le Polain de Waroux O, O’Brien KL, Edmunds WJ. The serotype distribution among healthy carriers before vaccination is essential for predicting the impact of pneumococcal conjugate vaccine on invasive disease. PLoS computational biology. 2015;11:e1004173.

[45] Adegbola RA, DeAntonio R, Hill PC, Roca A, Usuf E, Hoet B, et al. Carriage of Streptococcus pneumoniae and Other Respiratory Bacterial Pathogens in Low and Lower-Middle Income Countries: A Systematic Review and Meta-Analysis. PloS one. 2014;9:e103293.

[46] UNFPA. UNFPA State of the World Population 2014. 2015.

[47] Abdullahi O, Karani A, Tigoi CC, Mugo D, Kungu S, Wanjiru E, et al. Rates of acquisition and clearance of pneumococcal serotypes in the nasopharynges of children in Kilifi District, Kenya. The Journal of infectious diseases. 2012;206:1020–9.

[48] Melegaro A, Gay NJ, Medley GF. Estimating the transmission parameters of pneumococcal carriage in households. EpidemiolInfect. 2004;132:433–41.

[49] Nair H, Simoes EA, Rudan I, Gessner BD, Azziz-Baumgartner E, Zhang JS, et al. Global and regional burden of hospital admissions for severe acute lower respiratory infections in young children in 2010: a systematic analysis. Lancet. 2013;381:1380–90.

[50] Fleming-Dutra KE, Conklin L, Loo JD, Knoll MD, Park DE, Kirk J, et al. Systematic review of the effect of pneumococcal conjugate vaccine dosing schedules on vaccine-type nasopharyngeal carriage. The Pediatric infectious disease journal. 2014;33 Suppl 2:S152–60.

[51] Ota MO, Akinsola A, Townend J, Antonio M, Enwere G, Nsekpong D, et al. The immunogenicity and impact on nasopharyngeal carriage of fewer doses of conjugate pneumococcal vaccine immunization schedule. Vaccine. 2011;29:2999–3007.

